# DEAD-box RNA helicase 18 disrupts IRF3-binding to the interferon-β promoter

**DOI:** 10.1101/2021.10.26.465893

**Authors:** Xun Xiao, Mohan Wang, Wenkai Zhao, Puxian Fang, Yanrong Zhou, Dang Wang, Liurong Fang, Shaobo Xiao

## Abstract

The production of type I interferons (IFN-α/β) requires strict control to avoid excessive activation during viral infections. The binding of interferon regulatory factor 3 (IRF3) to the *IFN-β* promoter region in the nucleus is essential for IFN-β transcription; however, whether nuclear factors have important negative-regulatory roles in this process is largely unknown. By screening for IRF3-interacting partners in the nucleus, we identified DEAD-box RNA helicase 18 (DDX18) as an important negative regulator of intranuclear IRF3. Overexpression of DDX18 suppressed virus- and IRF3-induced IFN-β production, whereas knockdown of DDX18 expression or knockout of the *DDX18* gene had opposite effects. Mechanistically, DDX18 interacts with IRF3 and decreases the binding of IRF3 to the *IFN-β* promoter after viral infection. DDX18 knockdown mice (*Ddx18^+/-^*) further demonstrated that DDX18 suppressed antiviral innate immunity *in vivo*. Thus, despite many members of the DDX family act as important positive regulators in the cytoplasm, DDX18 plays a unique “braking” role in balancing virus-induced type I IFN production.

## Introduction

Type I interferon (IFN-α/β) responses constitute the first line of host defense against viral pathogens(Li *et al*, 2018; Wang & Fish, 2019). During viral infection, activation of innate antiviral immunity begins with the detection and recognition of pathogen-associated molecular patterns (PAMPs) via germline-encoded pattern-recognition receptors (PRRs), such as retinoic acid-inducible gene I (RIG-I)-like receptors (RLRs), Toll-like receptors (TLRs), and intracellular DNA sensors(Janeway & Medzhitov, 2002; Kato *et al*, 2006; Sun *et al*, 2013; Takeuchi & Akira, 2010; Yoneyama *et al*, 2004). Upon stimulation with viral PAMPs, these PRRs initiate signal transduction pathways that activate different key adaptors, including the caspase recruitment domain-containing mitochondrial antiviral signaling (MAVS)(Meylan *et al*, 2005; Seth *et al*, 2005), Toll/interleukin-1 receptor domain-containing adaptor that induces IFN-β (TRIF)(Oshiumi *et al*, 2003; Yamamoto *et al*, 2003), and stimulator of interferon genes (STING)(Ishikawa *et al*, 2009), the latter of which recruits TANK-binding kinase 1 (TBK1)/inhibitor-κB kinase ε (IKKε) kinases and results in phosphorylation of cytoplasmic interferon regulatory factor 3 (IRF3). After phosphorylation, IRF3 forms homodimers that move into the nucleus, where they bind to the promoters of type I IFN genes and lead to the production of type I IFNs and antiviral immune responses(Fitzgerald *et al*, 2003; Sharma *et al*, 2003). In this process, IRF3 acts as a master switch for the activation of *IFN-β* transcription(Sato *et al*, 2000).

Many positive regulators of IRF3 in the cytoplasm have been identified during viral infections. For example, RIO kinase 3 (RIOK3) bridges TBK1 and IRF3 to promote IFN-β production(Feng *et al*, 2014), and uiquitin-specific protease 22 (USP22) promotes the nuclear translocation of IRF3 and the expression of IFN-β by de-ubiquitinating and inhibiting the degradation of karyopherin α2 (KPNA2)(Cai *et al*, 2020). Although type I IFN production plays a critical role in host defense against viral invasion, uncontrolled type I IFN expression may lead to autoimmune diseases, such as systemic lupus erythematosus(Shrivastav & Niewold, 2013). Several important negative regulators have been found that degrade or dephosphorylate IRF3 and ensure that appropriate amounts of type I IFNs are produced during viral infection. For example, tripartite motif-containing protein (TRIM) 21 suppresses type I IFN production by driving the ubiquitination and degradation of activated IRF3(Yang *et al*, 2013). Protein phosphatase-2A (PP-2A) dephosphorylates activated IRF3, which prevents it from entering the nucleus and initiating the transcription of *IFN-β*(Long *et al*, 2014). Sentrin-specific protease 2 (SENP2) inhibits type I IFN production by deSUMOylating IRF3 and conditioning it for ubiquitination and degradation(Hu & Liao, 2017). In general, these reported regulators of IRF3 primarily target cytoplasmic IRF3. Many factors in the nucleus have been reported as regulators of type I IFN production, such as the transmembrane adaptor TRIM5α(Portilho *et al*, 2016) and interferon-inducible protein 16 (IFI16)(Ansari *et al*, 2015). However, whether these nuclear factors can specifically target IRF3, especially for negative regulation of IFN production, is largely unknown.

DEAD-box helicase 18 (DDX18) is a highly conserved member of the DDX helicase family in eukaryotes. In zebrafish, DDX18 was shown to be critical for cell cycle progression during primitive hematopoiesis(Payne *et al*, 2011). Its ortholog in yeast is HAS1, which has been reported to be essential for ribosome biogenesis(Dembowski *et al*, 2013). A recent study showed that DDX18 is located in the outer region of the nucleolus and plays an important role in early embryonic development and maintenance of pluripotency in embryonic stem cells(Zhang *et al*, 2020a). However, whether mammalian DDX18 is involved in regulation of innate antiviral immune responses remains unclear. In this study, we identified DDX18 as a novel negative regulator of nuclear IRF3 by screening for IRF3-interacting partners in the nucleus. DDX18 interacts with IRF3 via its helicase core domain, which results in a significant decrease in binding between IRF3 and the *IFN-β* gene promoter after Sendai-virus (SeV) infection. This novel function and mechanism of action for DDX18 give new insights into the poorly understood negative regulation of IRF3 in the nucleus.

## Results

### Identification of IRF3-bound protein in the nucleus

IRF3 is the master transcription factor for type I IFN production, and the binding of IRF3 to the *IFN-β* gene promoter in the nucleus is the last step for *IFN-β* transcription(Sato *et al*., 2000). Therefore, we screened for IRF3-interacting proteins in the nucleus to identify potential negative regulators of IRF3. To this end, HEK293T cells were transfected with an IRF3 expression construct and nuclear proteins were extracted by co-immunoprecipitation (Co-IP) with an antibody specific for human IRF3, followed by liquid chromatography-tandem mass spectrometry (LC-MS/MS; Fig. 1A). Some evident protein bands were observed after silver staining, and IRF3 expression was confirmed by western blot of cell lysates and Co-IP eluted samples from the IRF3-overexpression group compared with empty vector-transfected control cells (Fig. 1B). LC-MS/MS analysis showed a total of 66 differentially expressed IRF3-interacting proteins in the IRF3 overexpression group compared with the control group (listed in **Supplementary Table 1**).

**Figure 1.**
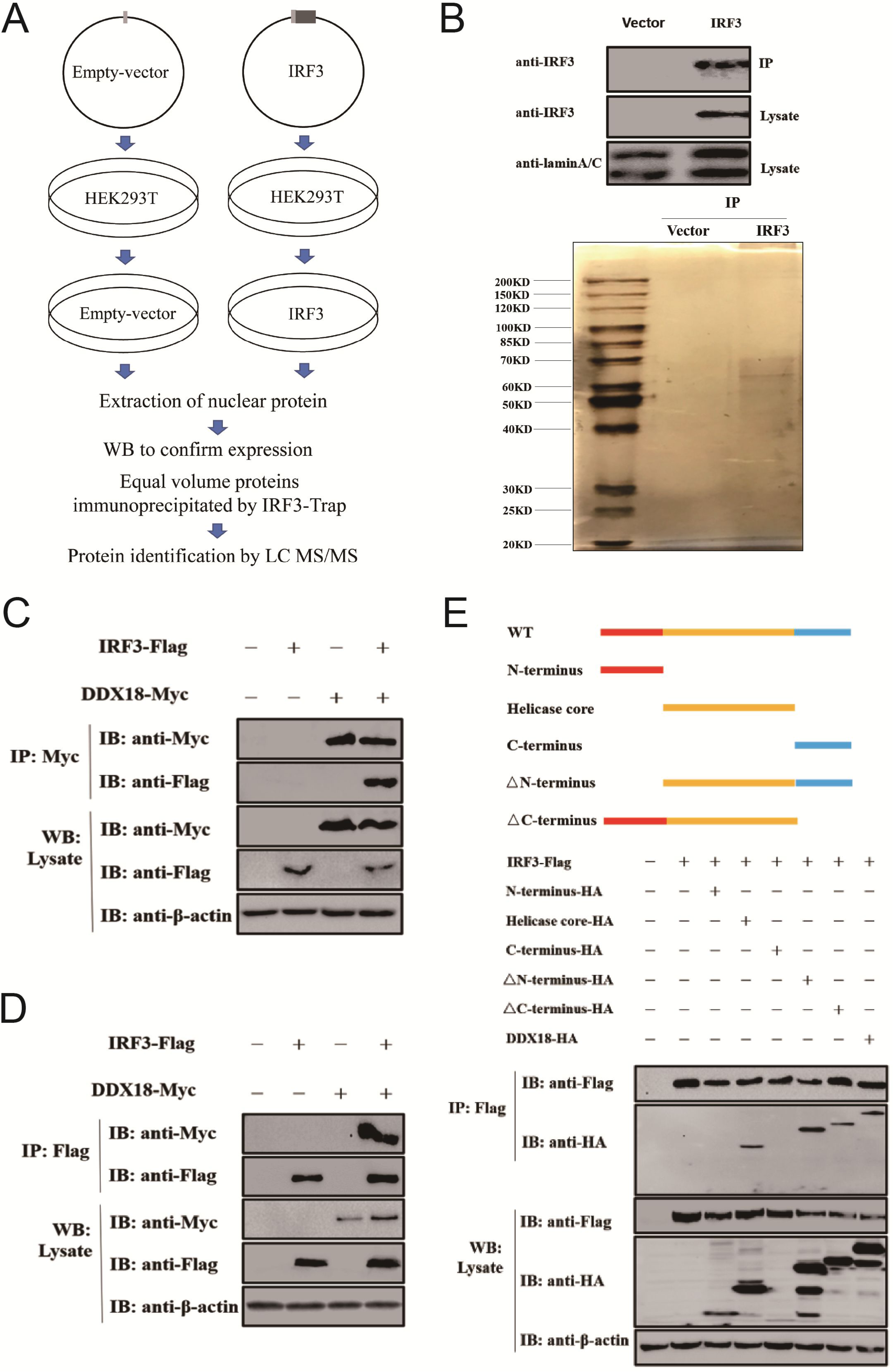
Screening of nuclear proteins interacting with IRF3. **A** Strategy used for screening interacting partners of IRF3 in the nucleus via co-immunoprecipitation (Co-IP) and liquid chromatography-tandem mass spectrometry (LC-MS/MS). **B** The extracted nuclear proteins (immunoprecipitated with anti-IRF3 beads) from empty vector and IRF3 overexpressed cells were detected by western blot (**top)** and a silver-stained SDS-PAGE analysis **(bottom)**. **C, D** HEK293T cells were co-transfected with Flag-tagged IRF3 and Myc-tagged hDDX18 expression plasmids. At 36 h post-transfection, Co-IP experiments were performed with anti-Flag antibody **(C)** and anti-Myc antibody. **D**, **E** Schematic diagram of full-length hDDX18 and its truncated mutants (**top**). HEK293T cells were co-transfected with expression constructs encoding HA-tagged hDDX18 or its truncated mutants together with Flag-tagged IRF3 expression plasmid. At 36 h post-transfection, Co-IP experiments were performed with anti-Flag antibody (**bottom**).

Among these proteins, DDX18 attracted the most attention. DDX18 belongs to the DDX helicase family and is mainly located in the nucleus(Zhang *et al*., 2020a). At present, DDX18 function is mainly confined to hematopoiesis and ribosome synthesis(Dembowski *et al*., 2013; Payne *et al*., 2011; Zhang *et al*., 2020a), and there has been no report regarding a potential role in the innate immune response. To further confirm the interaction between DDX18 and IRF3, we cloned the full-length cDNA of human DDX18 (hDDX18), and the expression construct encoding Myc-tagged hDDX18 was co-transfected with Flag-tagged IRF3 (human IRF3) into HEK293T cells, followed by immunoprecipitation assays with antibodies against Flag or Myc. Both the “forward and backward” pull-down assays supported an interaction between DDX18 and IRF3 (Fig. 1C, 1D). The N-terminal domain, the C-terminal disordered region/low complexity domain, and the helicase core region are three main functional domains of DDX18(Schütz *et al*, 2010). To investigate which DDX18 domain is required for the interaction between DDX18 and IRF3, five DDX18 mutants were constructed that included the N-terminus, helicase core, C-terminus, ΔN-terminus, and ΔC-terminus. As shown in Fig. 1E, overexpressed IRF3 immunoprecipitated with the helicase core, ΔN-terminus, and ΔC-terminus but not with the N-terminus or C-terminus of DDX18, suggesting that DDX18 interacts with IRF3 via its helicase core domain. These results impelled us to focus on DDX18 and further explore whether DDX18 plays a regulatory role in type I IFN production.

### DDX18 negatively regulates type I IFN production during viral infections

To determine whether DDX18 has an effect on type I IFN production, luciferase reporter assays were used to detect changes in transcriptional activity of the *IFN-β* promoter in cells overexpressing hDDX18. As shown in Fig. 2A, overexpression of hDDX18 in HEK293T significantly decreased the transcriptional activation of the *IFN-β* promoter induced by SeV, and the decrease was positively correlated with the level of hDDX18 expression. Consistently, significant reductions in the mRNA expression of *IFN-β*, interferon-stimulated gene (*ISG)15*, and *ISG56* induced by SeV were also observed in RAW264.7 cells infected with recombinant lentivirus expressing mouse DDX18 (mDDX18) (Fig. 2B). These results suggested DDX18 was a potent negative regulator of IFN-β production.

**Figure 2.**
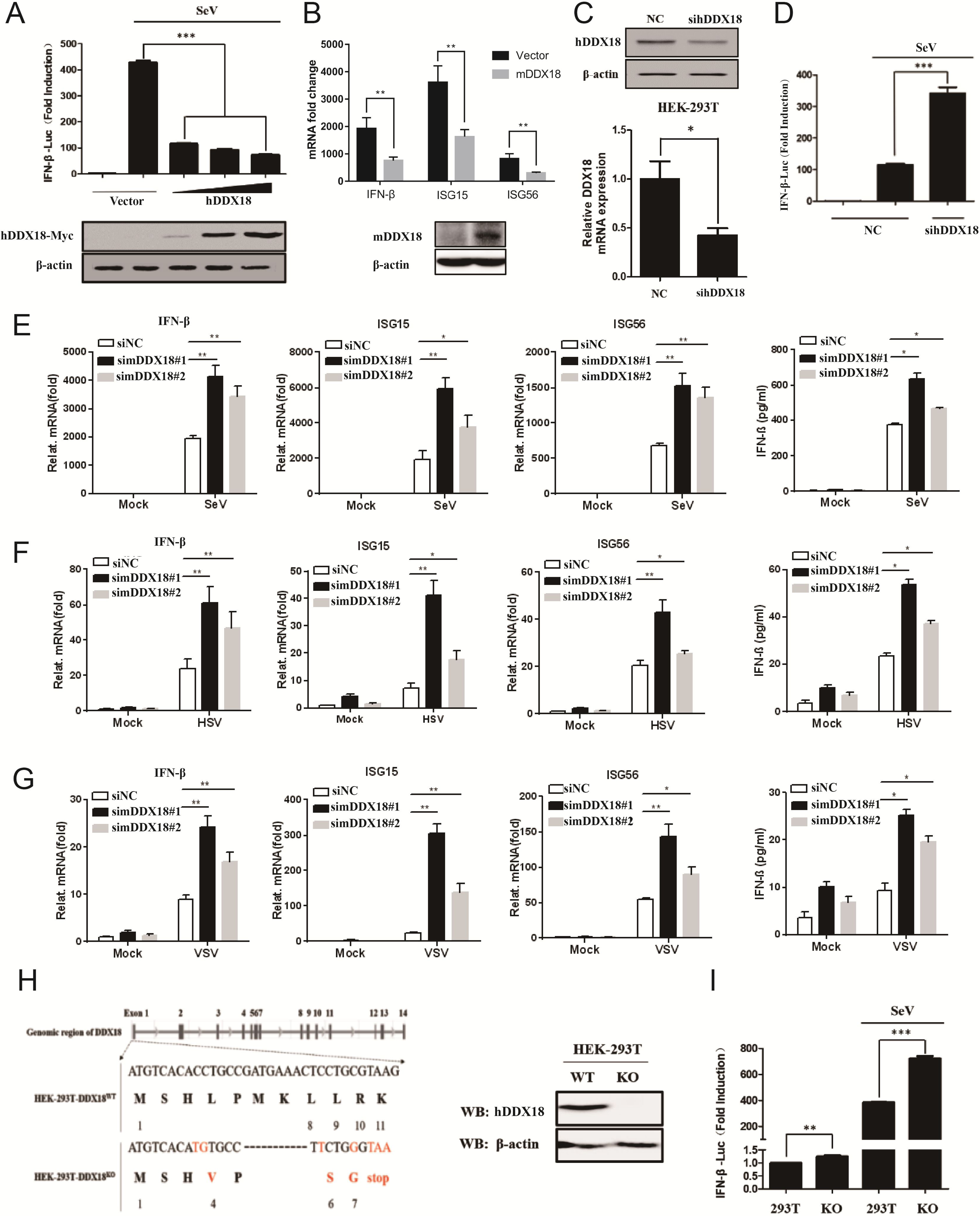
DDX18 negatively regulates virus-induced type I IFN production. **A** HEK293T cells were co-transfected with reporter plasmids IFN-β-Luc and pRL-TK, together with increasing amounts of hDDX18 expression plasmid or empty vector. At 24 h post-transfection, cells were mock-infected or infected with SeV (10 hemagglutination units/well). The cells were then subjected to dual-luciferase reporter assays at 12 h post-infection. **B** RAW264.7 cells were infected with recombinant lentivirus expressing the mouse *Ddx18* gene (mDDX18) or empty lentivirus, and then infected with SeV for 24 h. The mRNA levels of *Ifn-β*, *Isg15*, and *Isg56* were detected by qPCR. The mean mRNA level of the empty vector group without viral stimulation was defined as 1 standard unit. **C** HEK293T cells were transfected with hDDX18-specific (sihDDX18) or control (siNC) siRNAs, and the knockdown effects were analyzed by western blot and qPCR. **D** HEK293T cells were co-transfected with sihDDX18 or siNC and the reporter plasmids IFN-β-Luc and pRL-TK. The cells were mock-infected or infected with SeV (10 hemagglutination units/well). **E–G** RAW264.7 cells were transfected with mDDX18-specific (simDDX18) or control (siNC) siRNAs and mock-infected or infected with SeV **(E)**, HSV **(F),** or VSV **(G)** for another 24 h. The mRNA levels of *Ifn-β*, *Isg15*, and *Isg56* were detected by qPCR, and IFN-β protein levels in cell supernatants were detected by enzyme-linked immunosorbent assay (ELISA). **H** The effects of DDX18 knockout in HEK293T-DDX18-KO cells were analyzed by western blotting. **I** HEK293T or HEK293-DDX18-KO cells were transfected with the reporter plasmids IFN-β-Luc and pRL-TK. At 24 h post-transfection, the cells were mock-infected or infected with SeV. The data were obtained from three independent experiments (mean ± SD). *p < 0.05, **p < 0.01, ***p < 0.001.

To further confirm the negative regulatory function of DDX18 in IFN-β production, we used DDX18-specific siRNA to knockdown the expression of endogenous DDX18 (Fig. 2C) and demonstrated that knockdown of hDDX18 significantly enhanced SeV-induced *IFN-β* promoter activation (Fig. 2D) in HEK293T cells. Consistent with hDDX18, knockdown of mDDX18 significantly enhanced SeV-induced expression of *IFN-β*, *ISG15*, and *ISG56* mRNA as well as IFN-β secretion in RAW264.7 cells, and the degree of enhancement was associated with knockdown efficiency (Fig. 2E**, Supplementary Fig. 1)**. Similar trends were observed after infection with herpes simplex virus type 1 (HSV-1) (Fig. 2F) and vesicular stomatitis virus (VSV) (Fig. 2G).

In addition to overexpressing and interfering with DDX18 expression, we knocked out the *DDX18* gene in HEK293T cells using CRISPR/Cas9 technology (Fig. 2H) and determined that knockout of the *DDX18* gene significantly enhanced *IFN-β* promoter activity induced by SeV infection (Fig. 2I). Taken together, these data indicated that DDX18 contributed to a significant negative regulation of virus-induced IFN-β production and ISG expression.

### DDX18 inhibits type I IFN production by targeting intranuclear IRF3

Previous study showed that many DDX members located in the nucleus have been reported to migrate into the cytoplasm and target key cytoplasmic regulatory factors that mediate innate immune responses during viral infection(Wu *et al*, 2021). Therefore, we performed luciferase reporter assays to examine the effects of DDX18 on *IFN-β* promoter activation mediated by various adaptors/kinases and confirm whether DDX18 negatively regulated type I IFN production by targeting nuclear IRF3. As shown in Fig. 3A, overexpression of hDDX18 strongly reduced *IFN-β* promoter activation induced by RIG-I, RIG-IN, MDA5, IPS-1, TBK1, IKKε, IRF3, and IRF3-5D (a constitutively active form of IRF3), which suggested that hDDX18 functioned downstream of IRF3. The decrease in promoter activation was positively correlated with the increased expression levels of hDDX18 (Fig. 3B). However, stronger *IFN-β* promoter activation was observed after overexpression of IRF3 in both the hDDX18-knockdown and -knockout cells (Fig. 3C, 3D). Together, these data indicated that DDX18 indeed negatively regulated the production of type I IFN by targeting IRF3 in the nucleus.

**Figure 3.**
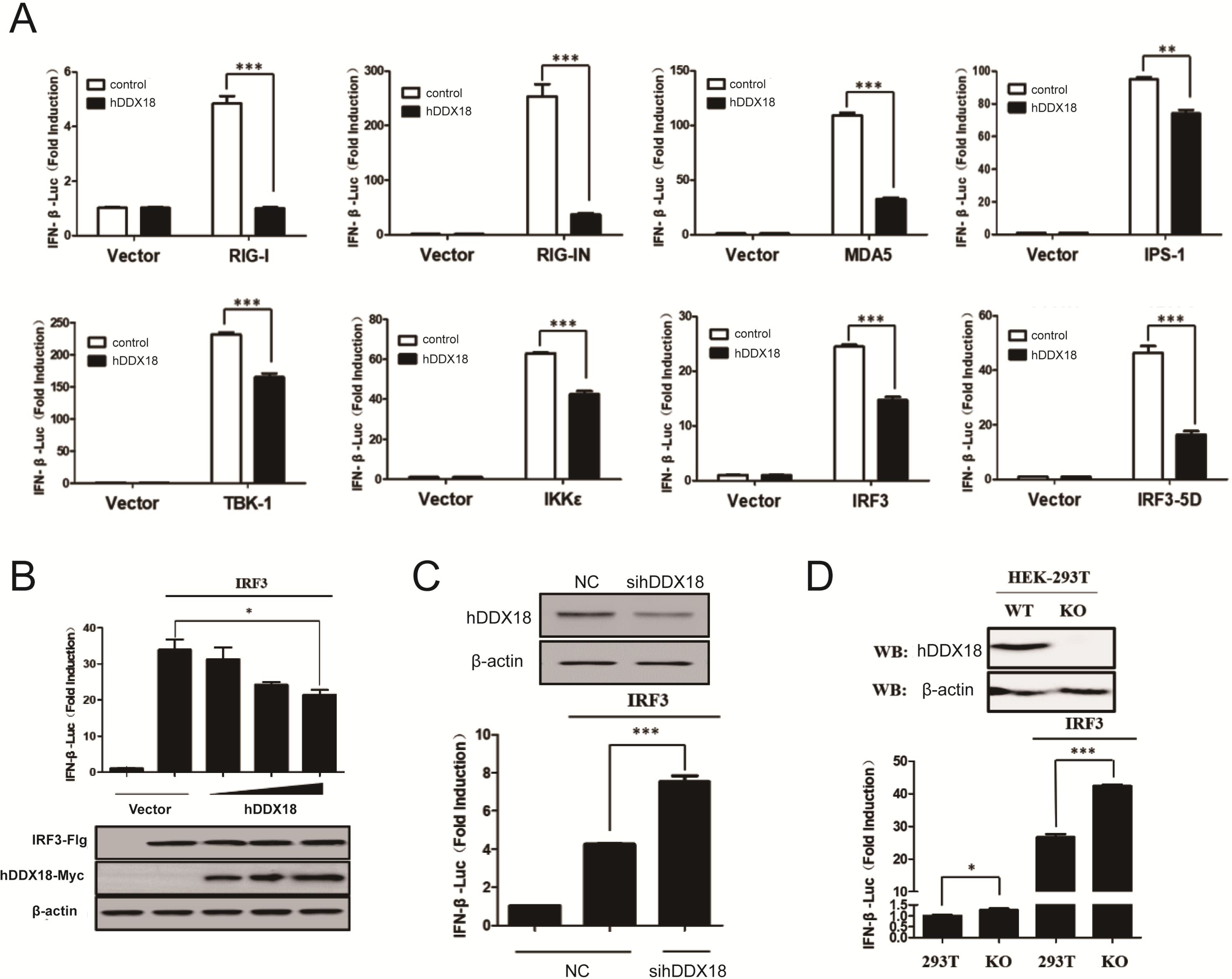
DDX18 inhibits type I IFN production by targeting intranuclear IRF3. **A** HEK293T cells were co-transfected with the reporter plasmids IFN-β-Luc, pRL-TK, and hDDX18 expression plasmid or empty vector along with plasmids expressing RIG-I, RIG-IN, MDA5, IPS-1, TBK1, IKKε, IRF3, or IRF3-5D. At 48 h after co-transfection, the cells were collected for dual-luciferase reporter assays. **B** HEK293T cells were co-transfected with IRF3 expression plasmid, IFN-β-Luc, and pRL-TK plasmids, along with increasing amounts of the expression construct encoding hDDX18. **C** HEK293T cells were co-transfected with IFN-β-Luc, pRL-TK, and IRF3 expression plasmids, along with sihDDX18 or siNC. **D** HEK293-DDX18-KO cells were co-transfected with IFN-β-Luc and pRL-TK plasmid, along with the IRF3 expression plasmid. At 48 h after co-transfection, cells were collected for dual-luciferase reporter assays. The data were obtained from three independent experiments (mean ± SD). *p < 0.05, **p < 0.01, ***p < 0.001.

### ATPase hydrolytic activity and RNA-binding activity of DDX18 are not required for suppression of type I IFN production

DDX proteins belong to the family of ATPase-dependent RNA helicases, and the helicase core domain possesses two main functions: ATPase hydrolytic activity and RNA-binding activity(Collins *et al*, 2009; Hodge *et al*, 2011). These two functions are necessary for some members of the DDX family to negatively regulate type I IFN production by competing with RNA substrates, and DDX18 interacts with IRF3 via its helicase core domain (Fig. 1E). Therefore, we asked whether DDX18 is dependent on these two helicase activities for inhibition of type I IFN production. A previous study reported that switching from K to A or E in Motif I of the DEAD-box family results in loss of ATPase hydrolytic activity, while switching from S to L in motif III results in loss of RNA-binding activity(Li *et al*, 2014). Thus, we generated a total of four DDX18 mutants, including hDDX18 K229A and S364L and mDDX18 K219E and S354L **(Supplementary Fig. 2A and 2B)**. Specifically, the hDDX18 mutations K229A and S364L abolished ATPase hydrolytic activity and RNA-binding ability, respectively; the mDDX18 mutations K219E and S354L abolished ATPase hydrolytic activity and RNA-binding ability, respectively. Luciferase assays showed that hDDX18 and its two mutants exerted comparable inhibitory effects on the activation of the *IFN-β* promoter induced by SeV (Fig. 4A) and IRF3 (Fig. 4B). Furthermore, overexpression of hDDX18 and its two mutants in DDX18-knockout HEK293T cells restored activation of the *IFN-β* promoter induced by SeV (Fig. 4C) and IRF3 (Fig. 4D). We did not observe significant differences in inhibition of type I IFN production between hDDX18 and its mutants. Significant reductions in *IFN-β*, *ISG15*, and *ISG56* mRNA induced by SeV (Fig. 4E) and VSV (Fig. 4F) were observed in RAW264.7 cells infected with recombinant lentivirus expressing mDDX18 or its two mutants. Together, these data indicated that neither ATPase hydrolytic activity nor RNA-binding activity of DDX18 is required for its negative regulatory effect on type I IFN production. Hence, DDX18 may directly bind to IRF3 to disable intranuclear IRF3 activity.

**Figure 4.**
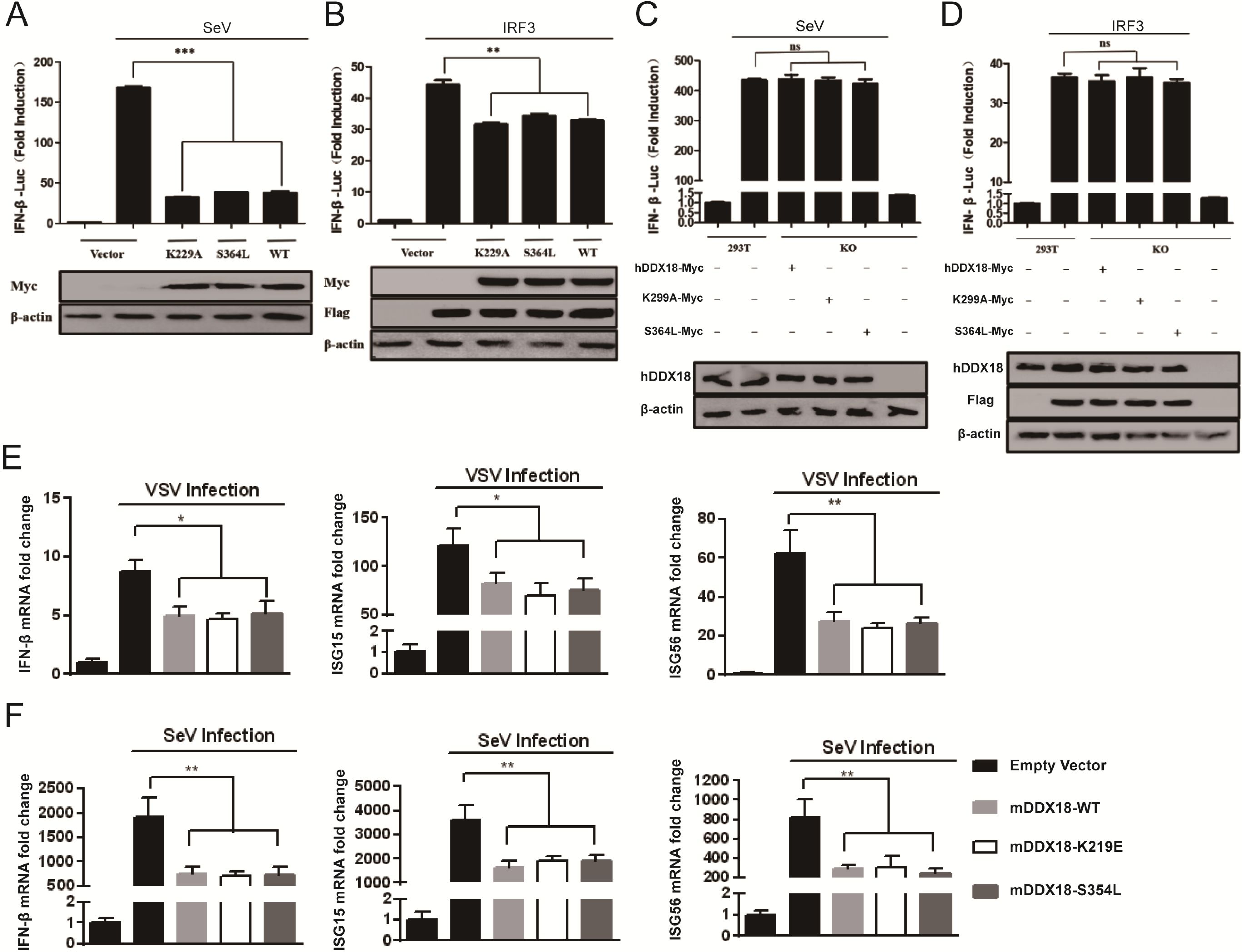
Neither ATPase hydrolytic activity nor RNA-binding activity of DDX18 is required for suppression of type I IFN production. **A, C** HEK293T or HEK293T-DDX18-KO cells were co-transfected with the reporter plasmids IFN-β-Luc and pRL-TK and plasmids expressing hDDX18 (WT), hDDX18 mutants (K229A, S364L), or empty vector. At 24 h post-transfection, the cells were mock-infected or infected with SeV, followed by dual-luciferase assays at 12 h post-infection. **B, D** HEK293T cells and HEK293T-DDX18-KO cells were co-transfected with IRF3 expression plasmid, IFN-β-Luc and pRL-TK, and expression plasmids encoding hDDX18 (WT) or hDDX18 mutants (K229A, S364L). At 48 h after co-transfection, cells were collected for dual-luciferase reporter assays. **E, F** RAW264.7 cells were infected with recombinant lentivirus expressing mDDX18, its mutants (mDDX18-K219E, mDDX18-S354L), or empty vector, and then infected with VSV **(E)** or SeV **(F)** for another 24 h. The mRNA levels of *Ifn-β*, *Isg15*, and *Isg56* were detected by qPCR. All data were obtained from three independent experiments (mean ± SD). *p < 0.05, **p < 0.01, ***p < 0.001.

### DDX18 reduces IRF3-binding to the *IFN-β* promoter

The last step of RLR-mediated type I IFN transcription is the binding of phosphorylated IRF3 dimers to the *IFN-β* promoter in the nucleus(Sato *et al*., 2000). We hypothesized that the interaction between DDX18 and intranuclear IRF3 would result in reduction of the binding between IRF3 and the *IFN-β* promoter. To this end, chromatin immunoprecipitation (ChIP) assays were performed in SeV-infected HEK293T cells overexpressing hDDX18, hDDX18 mutant S364L, or empty vector. The qPCR results showed that *IFN-β* promoter DNA immunoprecipitated with histone 3 (positive control) but not with hDDX18 or its mutant (Fig. 5A), which suggested that there was no interaction between hDDX18 and the *IFN-β* promoter. However, after SeV infection, overexpression of hDDX18 or its mutant significantly reduced the level of *IFN-β* promoter DNA interacting with IRF3 but not with histone 3 (Fig. 5B). These data indicated that DDX18 specifically interfered with the DNA-binding ability of IRF3 in the nucleus. Taken together, we concluded that DDX18 interacted with the activated IRF3 entering into the nucleus and disrupted the binding between IRF3 and the *IFN-β* promoter, thus, efficiently inhibiting the production of type I IFN triggered by viral infection.

**Figure 5.**
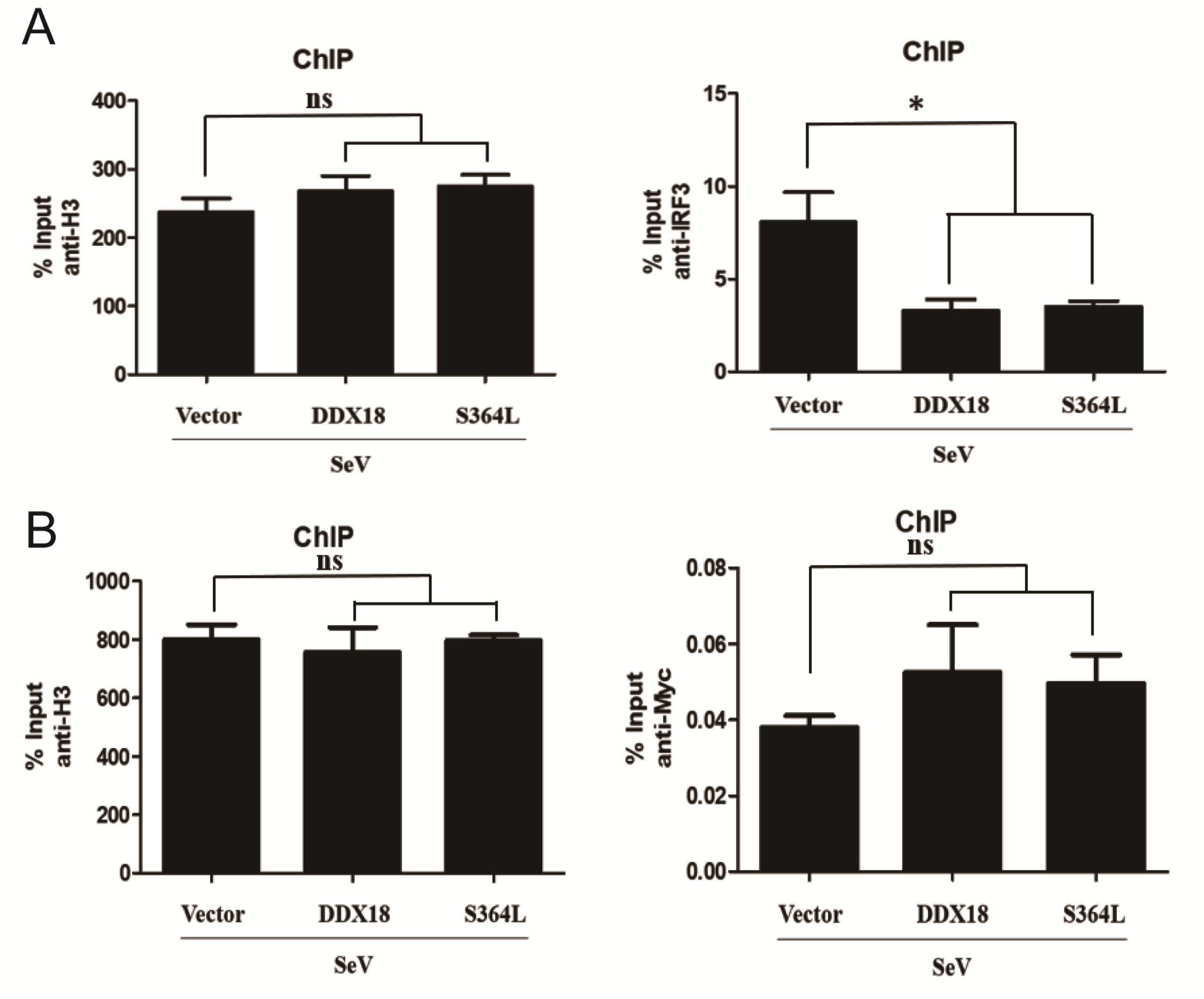
DDX18 reduces IRF3-binding to the *IFN-β* promoter. **A, B** HEK293T cells were transfected with expression plasmids containing Myc-DDX18 or its mutant Myc-DDX18-S364L. At 24 h post-transfection, the cells were infected with SeV followed by chromatin immunoprecipitation (ChIP) analysis with anti-Myc **(A)** or anti-IRF3 **(B)** antibody. Anti-histone 3 was used as a positive control. All data were obtained from three independent experiments (mean ± SD). *p < 0.05, **p < 0.01, ***p < 0.001.

### *In vivo* knockdown of DDX18 enhances antiviral responses

To further investigate the function of DDX18 *in vivo*, we used CRISPR/Cas9 knockout technology to create targeted deletion of the *Ddx18* gene in mice (**Supplementary Fig. 3A**). A large number of heterozygous mice (*Ddx18^+/-^*) and wild-type mice (*Ddx18^+/+^*) were obtained but not homozygous DDX18 knockout mice (*Ddx18^-/-^*), suggesting that the *Ddx18* gene may play an indispensable role in mouse development (**Supplementary Fig. 3B**). Western blot analysis demonstrated that the expression of DDX18 in *Ddx18^+/-^* mice was lower than that in *Ddx18^+/+^* mice (**Supplementary Fig. 3C**). However, *Ddx18^+/-^* and *Ddx18^+/+^* mice were similar in body type, weight, and fertility. We used *Ddx18^+/-^* and *Ddx18^+/+^* mice to study the function of DDX18 *in vivo*. The expression of DDX18 protein in macrophages from *Ddx18^+/-^* mice was significantly lower than in macrophages from wild-type littermates (**Supplementary Fig. 4A, 4B**). After infections with SeV, VSV, and HSV-1, the expression of *Ifn-β*, *Isg15*, and *Isg56* mRNA was significantly higher in macrophages from *Ddx18^+/-^* mice than in macrophages from wild-type littermates (**Supplementary Fig. 4C–E**).

The *in vivo* ability of DDX18 to inhibit type I IFN production was mainly determined by mouse survival, tissue and serum IFN levels, tissue viral loads, and organ injury. As shown in Fig. 6A, the survival rate of *Ddx18^+/-^* mice was higher than wild-type mice after tail vein injection of HSV-1, and the time until death of the *Ddx18^+/-^* mice was obviously delayed. Improvement in survival was associated with decreased viral loads in the brain tissue of infected *Ddx18^+/-^* mice (Fig. 6B). Hematoxylin and eosin staining showed that the degree of alveolar interstitial congestion, edema, and inflammatory cell infiltration in the lung tissue of *Ddx18^+/-^* mice was milder than that of wild-type mice (Fig. 6C). The qPCR results revealed that the mRNA levels of *Ifn-β*, *Isg15*, and *Isg56* in the brain and lung tissue of *Ddx18^+/-^* mice were obviously higher than in wild-type mice (Fig. 6D **and** 6E), and the average level of IFN-β in serum from *Ddx18*^+/-^ mice was > 2-fold higher than in wild-type mice (Fig. 6F). In addition to HSV-1, we examined the *in vivo* effect of DDX18 during VSV infection. A lower viral burden (Fig. 7A, 7B) and greater IFN-β production and *Ifn-β*, *Isg15*, and *Isg56* mRNA expression (Fig. 7C–F) were observed in *Ddx18^+/-^* mice compared with wild-type littermates after VSV infection. These data demonstrated convincingly that downregulation of DDX18 enhanced the natural antiviral immune response by promoting type I IFN production *in vivo*.

**Figure 6.**
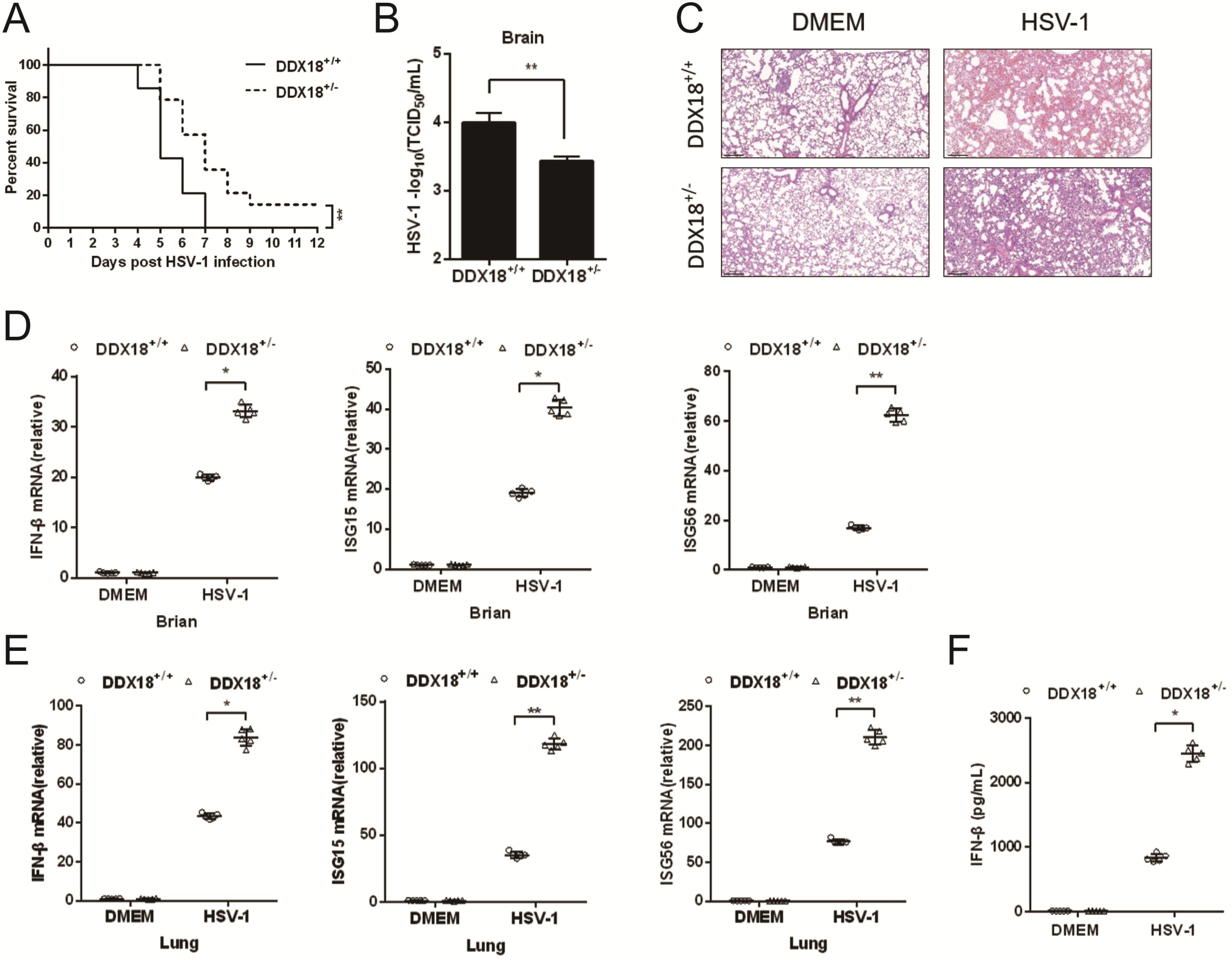
*In vivo* knockdown of DDX18 enhances the innate antiviral response against HSV-1 infection. **A** Survival rates for *Ddx18*^+/-^ and *Ddx18^+/+^* mice injected with HSV-1 (2.5 × 10^7^ PFU/mouse) via the tail vein. **B** Determination of viral loads in the brains of *Ddx18^+/-^* and *Ddx18^+/+^* mice after HSV-1 infection. **C** Hematoxylin and eosin staining of lung sections from *Ddx18^+/+^* or *Ddx18^+/-^* mice that were mock-infected or infected with HSV-1. **D, E** qRT-PCR analysis of *Ifn-β*, *Isg15*, and *Isg56* mRNA in the brains **(D)** and lungs **(E)** of *Ddx18^+/+^* or *Ddx18^+/-^* mice at 72 h after intravenous injection of DMEM medium (mock-infected) or HSV-1. **F** Detection of IFN-β protein levels in sera from HSV-1-infected *Ddx18^+/+^* or *Ddx18^+/-^* mice determined by ELISA. *p < 0.05, **p < 0.01, ***p < 0.001.

**Figure 7.**
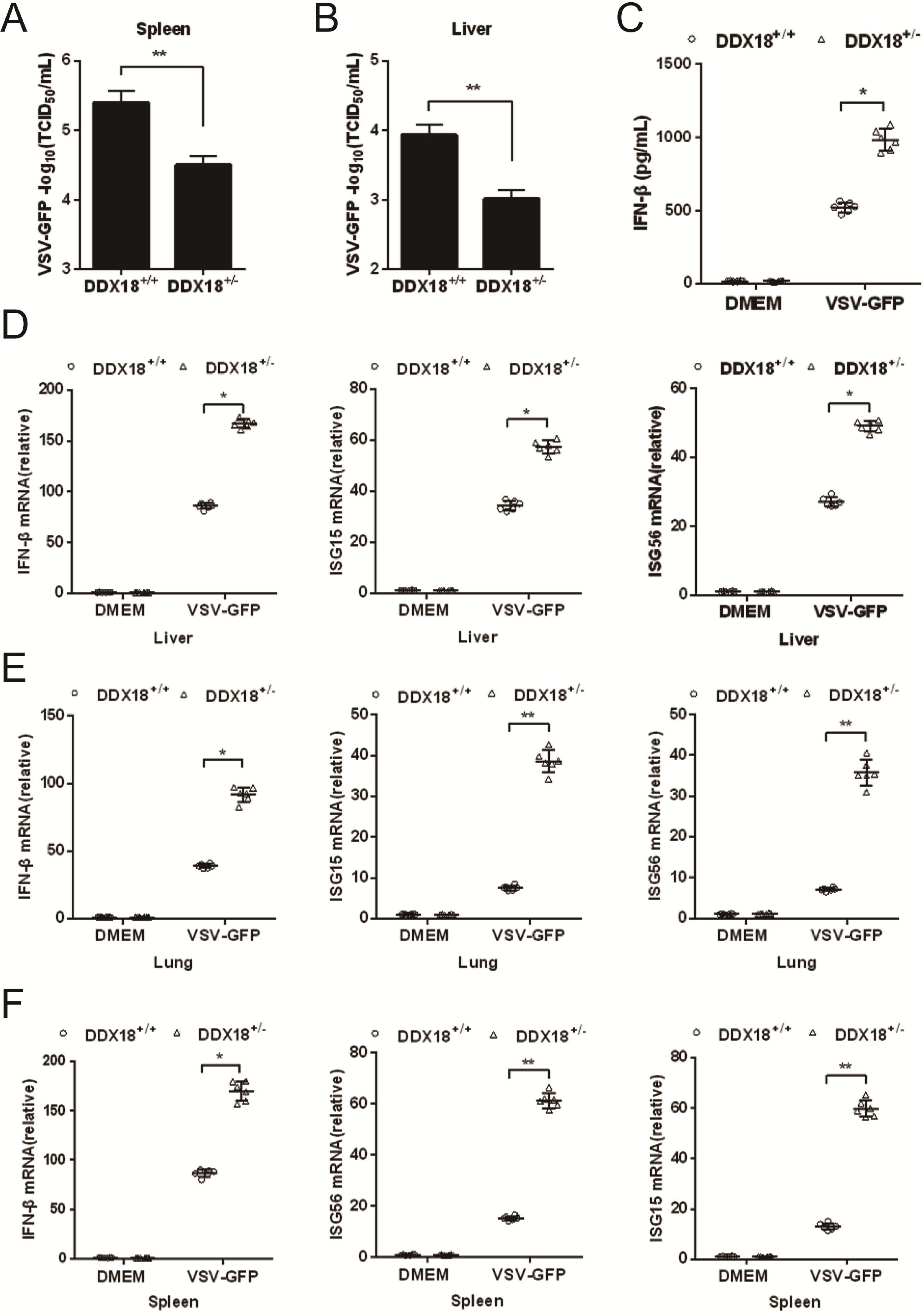
*In vivo* knockdown of DDX18 enhances the innate antiviral response against VSV infection. **A, B** Virus loads in spleens (**A**) or livers (**B**) from *Ddx18^+/-^* and *Ddx18^+/+^* mice after injection of VSV (1 × 10^9^ PFU/mouse) via the tail vein. **C** Detection of IFN-β protein levels in sera from VSV-infected *Ddx18^+/-^* or *Ddx18^+/+^* mice determined by ELISA. **D–F** qRT-PCR analysis of *Ifn-β*, *Isg15*, and *Isg56* mRNA in livers **(D),** lungs **(E),** and spleens **(F)** from VSV-infected *Ddx18*^+/+^ or *Ddx18*^+/-^ mice. *p < 0.05, **p < 0.01, ***p < 0.001.

## Discussion

As a class of ATPase activity-dependent RNA helicases, DDX family members are highly conserved in evolution, widely distributed in eukaryotes, and play a critical role in transcription, translation, ribosome biogenesis, and RNA transportation. Progressively more evidence has indicated that, in addition to their traditional functions in RNA metabolism, several DDX proteins are closely related to innate immune responses(Taschuk & Cherry, 2020). For example, DDX3(Valiente-Echeverría *et al*, 2015), DDX21(Zhang *et al*, 2011a), DDX23(Ruan *et al*, 2019), and DDX58(Loo & Gale, 2011; Yoneyama *et al*., 2004) have been reported as additional receptors for viral dsRNA that positively regulate RLR-mediated type I IFN production; DDX41 senses dsDNA from DNA viruses and upregulates type I IFN generation(Parvatiyar *et al*, 2012; Zhang *et al*, 2011b). Furthermore, studies in recent years have discovered that DDX19 and DDX46 can negatively regulate the production of type I IFN after viral infection(Zhang *et al*, 2019; Zheng *et al*, 2017). However, except for the nuclear protein DDX46, the reported DDX family members involved in regulating type I IFN response function mainly in the cytoplasm, and the majority are positive regulators. In this study, we used IRF3 to screen for IFN regulators in the nucleus and found that DDX18 interacts with IRF3. The results from *in vitro* and *in vivo* analyses clearly show that DDX18 is a negative regulator of type I IFN production after viral infection. To our knowledge, DDX18 is the second DDX member to be identified that functions in the nucleus to negatively regulate antiviral innate immunity.

After viral infection, DDX46 inhibits the production of type I IFN by entrapping m^6^A-demethylated antiviral transcripts in the nucleus(Zheng *et al*., 2017). This negative regulatory function of DDX46 is closely related to the RNA binding activity of DDX46. However, we found that neither ATPase hydrolytic nor RNA-binding activity of DDX18 was required for its suppression of type I IFN production. In addition, DDX46 functions downstream of MAVS and upstream of TBK1 while DDX18 and its mutants inhibited the production of type I IFN induced by various adaptors and transcription factors including IRF3, which suggested that DDX18 functioned downstream of IRF3. Therefore, despite their similarities, DDX18 and DDX46 act by completely different mechanisms to suppress type I IFN production. Another negative regulatory DDX family member, DDX19, has many similarities with DDX18. For example, both DDX18 and DDX19 interact with IRF3 through the helicase core region to disrupt IRF3 signaling functions, and neither of these proteins rely on ATPase hydrolytic or RNA-binding activity to suppress type I IFN production. However, DDX18 targets IRF3 in the nucleus, while DDX19 targets IRF3 in the cytoplasm. This difference creates two different stages for DDX18 and DDX19 negative regulatory function. Together, the diverse regulatory mechanisms of DDX18, DDX19, and DDX46 indicate that negative regulation of innate antiviral immunity by DDX family proteins is ideal and effective. When the immune system detects excessive IFN production, DDX46 can reduce the expression of some key antiviral signaling molecules, and DDX19 can inhibit the phosphorylation of IRF3. Both of these effects reduce the levels of activated IRF3 in the cytoplasm, while DDX18 acts as the final brake by preventing IRF3 from binding to the IFN promoter in the nucleus. These rich negative regulatory measures can ensure that the inhibition of excessive type I IFN production is maximized.

In this study, we used CRISPR/Cas9 knockout technology to create targeted deletion of *Ddx18* in mice to investigate the function of DDX18 *in vivo*. However, the *Ddx18^-/-^* mice were unable to survive, which is consistent with a recent report by Zhang *et al*. who demonstrated that knockout of DDX18 leads to embryonic lethality(Zhang *et al*, 2020b). Interestingly, similar to *Ddx18^-/-^* mice, homozygous *Ddx46^-/-^* and *Ddx19^-/-^* mice are not viable. Both DDX46 and DDX19 are negative regulators of type I IFN production, and it is likely that these DDX proteins are also essential for embryonic development. The DDX18, DDX19, and DDX46 deficient heterozygous mice (+/-) from this and other studies showed higher efficiency of viral clearance and produced larger amounts of type I IFN compared with the wild-type mice during viral infections. Considering that DDX family proteins are widely expressed in eukaryotic cells, these *in vivo* results strongly suggest that normal organisms may regulate the expression of these DDX family members to efficiently maintain the balance of type I IFNs in the body.

The question of how DDX18 affects the binding of IRF3 to the *IFN-β* promoter has not been fully resolved in this study. Previous studies have reported several cellular proteins that target the transcriptionally active form of IRF3 in the nucleus to inhibit its function. For example, TRIM26 mediates IRF3 ubiquitination and degradation in the nucleus to terminate IRF3 activation(Wang *et al*, 2015). However, we did not find that DDX18 had any effect on the expression of IRF3 after viral infections, which implies that DDX18 can neither degrade IRF3 protein nor retain *IRF3* mRNA in the nucleus to reduce IRF3 expression. We speculated that DDX18 may bind to IRF3 and recruit an important protease into the nucleus to increase or destroy modifications that affect IRF3 binding to DNA. At present, several negative regulatory proteins that can modify IRF3 have been reported, such as KAT8 and OTUD1(Huai *et al*, 2019; Zhang *et al*., 2020b). KAT8 directly interacts with IRF3 and mediates the acetylation of the K359 amino acid in the IRF3 protein(Huai *et al*., 2019), while OTUD1 cleaves the virus-induced K6-linked ubiquitination of IRF3(Zhang *et al*., 2020b). In both cases, the binding of IRF3 to the *IFN-β* promoter is significantly reduced. Similarly, our results showed that DDX18 interacts with IRF3 through the helicase core region, and the binding of IRF3 to the *IFN-β* promoter is significantly reduced in cells over-expressing DDX18 or its mutants. However, exactly which proteins DDX18 may recruit to influence the DNA binding activity of IRF3 remains unclear. Future studies will focus on proteins recruited by DDX18 that may interact with IRF3 to better understand this newly discovered cellular mechanism for down-regulating IRF3 activity.

In summary, our study demonstrated that the nuclear DDX protein DDX18 significantly inhibited the production of type I IFN after viral infection. Unlike the previously reported nuclear protein DDX46, DDX18 negatively regulates the antiviral innate response by targeting activated IRF3 rather than the mRNA of key antiviral molecules. The mechanism by which DDX18 acts on nuclear IRF3 is distinct from other known IRF3 inhibitors, and further investigations will be required to elucidate the mechanisms of interaction between DDX18 and nuclear IRF3 in response to viral infection.

## Materials and Methods

### Mice

C57BL6/J mice (6–8 weeks old) were obtained from the experimental animal center of Huazhong Agricultural University (Wuhan, China). *Ddx18*^+/−^ C57BL6/J mice were generated by CRISPR/Cas9-mediated genome editing and provided by Cyagen Biosciences Inc. (Guangzhou, China). The mice were housed in a specific pathogen free environment and challenged with SeV and sampled in a designated biosafety cabinet. All animal experiments were conducted in accordance with the guidelines of the Laboratory Animal Center of Huazhong Agricultural University. The protocols were approved by the Ethical Committee on Animal Research at Huazhong Agricultural University.

### Cells and viruses

HEK293T and RAW264.7 cells were obtained from the China Center for Type Culture Collection (Wuhan, China) and cultured in Dulbecco’s modified Eagle’s medium (DMEM; Invitrogen-Thermo Fisher; Waltham, MA, USA) supplemented with 10% fetal bovine serum (FBS) at 37 °C in a humidified atmosphere with 5% CO_2._ Mouse peritoneal macrophages (MPMs) were isolated from mice 4 d after the intraperitoneal injection of thioglycollate (Merck & Co.; Kenilworth, NJ, USA) as described previously(Zhang *et al*., 2019). MPMs were cultured in RPMI 1640 medium supplemented with 10% FBS at 37 °C with 5% CO_2_. SeV was purchased from the China Virus Resources and Information Center, Wuhan Institute of Virology, Chinese Academy of Sciences. HSV-1 strain F^45^ was a gift from Dr. Chunfu Zheng at Fujian Medical University, and VSV was a gift from Dr. Zhigao Bu at Harbin Veterinary Research Institute of the Chinese Academy of Agricultural Sciences.

### Plasmid constructions and RNA interference

The hDDX18, IRF3 and mDDX18 expression plasmids were constructed by RT-PCR amplification of cDNAs from HEK293T or RAW264.7 cells that were then cloned into pCAGGS-Flag, pCAGGS-Myc, or pCAGGS-HA vectors (with N-terminal Flag, Myc, or HA tags, respectively). All DDX18 substitution and deletion mutants were cloned into the pCAGGS vectors, and mutations were confirmed by DNA sequencing. The siRNAs targeting DDX18 or the negative control siRNA (Invitrogen) were transfected at a working concentration of 50 nM. The sequences of siRNAs and primers for plasmid construction can be provided upon request.

### Establishing DDX18 knockout cell lines

CRISPR/Cas9 technology was used to construct DDX18 knockout (KO) cell lines. The sgRNAs were designed using the http://tools.genome-engineering.org website and cloned into the PX459 vector. The recombinant plasmid was transfected into HEK293T cells, and the cells were screened for puromycin-resistance. The puromycin-resistant clones were propagated from single cells, and knockout cell lines were verified by DNA sequencing and western blot analysis using an anti-DDX18 antibody.

### Viral infection

HEK293T and RAW264.7 cells were infected with SeV (10 hemagglutinating activity units/well) for the indicated number of hours. RAW264.7 cells were infected with VSV (multiplicity of infection [MOI] = 10) or HSV-1 (MOI = 10) for the indicated times. MPMs from *Ddx18*^+/+^ or *Ddx18^+/-^* mice were infected with SeV (10 hemagglutinating activity units/well), HSV-1 (MOI = 10), or VSV (MOI = 10) for 12 h. For mouse infections, six to eight-week-old, sex-matched *Ddx18*^+/+^ or *Ddx18^+/-^* mice were infected intravenously with HSV-1 (2.5 × 10^7^ plaque-forming units [PFU] per mouse) and survival of the animals was monitored daily. The sera or tissues from mice infected with HSV-1 were collected at 18 h or 72 h post-infection, respectively, for enzyme-linked immunosorbent assays (ELISAs), qPCR, 50% tissue culture infectious dose (TCID_50_) assay, or histological analyses. For VSV infection, 2 × 10^9^ PFU per mouse were intravenously injected and the sera or tissues were collected at 18 h or 24 h post-infection, respectively, for ELISA, qPCR, or TCID50 assays.

### Lentiviral packaging

Recombinant lentiviruses expressing mDDX18 or its mutants were constructed and packaged as previously described(Ke *et al*, 2017). Briefly, lentiviral expression plasmids pLVX-IRES-mCherry-mDDX18, pLVX-IRES-mCherry-mDDX18-K219E, pLVX-IRES-mCherry-mDDX18-S354L were generated by cloning the cDNA of mDDX18 or its mutants into the lentiviral vector pLVX-IRES-mCherry (TaKaRa, Japan). To package the recombinant lentiviruses, lentiviral expression plasmids were co-transfected with the lentivirus-assisted plasmids pTRIP-VSV-G and pTRIP-Gag-Pol into HEK293T cells using Lipofectamine^®^ 3000 (Invitrogen). The supernatants were harvested at 48 h after co-transfection, followed by high-speed centrifugation (13,000 × g) for 4 h. The pellets were suspended in serum-free DMEM and stored at −70 °C until use.

### Dual-luciferase reporter assay

Cells were grown in 24-well plates to 50%–60% confluence and transfected in accordance with the Lipofectamine^®^ 2000 protocol. For luciferase reporter assays, the cells were co-transfected with the IFN-β-Luc reporter plasmid (0.1 μg/well), internal control plasmid pRL-TK (0.02 μg/well; Promega Corp.; Fitchburg, WI, USA), and the indicated expression constructs. In some cases, the cells were mock-infected or infected with SeV (10 hemagglutinating activity units/well) for 12 h. Subsequently, firefly and *Renilla reniformis* luciferase activities from lysed cells were determined using the dual-luciferase reporter assay system in accordance with the manufacturer’s protocol (Promega). Data are expressed as the relative firefly luciferase activities normalized to the activity of the *R. reniformis* luciferase.

### RNA extraction and quantitative real-time PCR (qRT-PCR)

The RNA extraction and qRT-PCR have been described previously(Ke *et al*, 2019). In brief, total RNA from cultured cells was extracted using TRIzol reagent (Omega Bio-Tek, Norcross, GA, USA) and reverse transcribed to cDNA using a Transcriptor First Strand cDNA Synthesis Kit (Roche, Mannheim, Germany) according to the manufacturer’s instructions. The qRT-PCR was performed with SYBR Green Real-Time PCR Master Mixes (Applied Biosystems, Foster City, CA, USA) using the Applied Biosystems ViiA 7 Real-Time PCR instrument. The mRNA levels of target genes were normalized to the expression level of the glyceraldehyde-3-phosphate dehydrogenase (GAPDH) gene. The GAPDH mRNA served as an internal reference.

### Nuclear and cytoplasmic protein extraction

The extraction of nuclear and cytoplasmic proteins was performed with the NE-PER Nuclear and Cytoplasmic Extraction Reagents (Thermo Fisher) in accordance with the manufacturer’s instructions. The purity of each fraction was determined according to the instructions in the kit.

### Co-IP and western blot

Co-IP and western blot analyses were performed as previously described(Tao *et al*, 2018). For Co-IP, the treated adherent cells were washed with phosphate-buffered saline (PBS) twice while in 60 mm cell culture dishes, followed by the addition of 1 mL of lysis buffer. The cell supernatants were collected by centrifugation, and a portion of supernatants were used in the whole cell extract assays. The remaining portion of supernatant was incubated with specific antibodies overnight at 4 °C and then treated with protein A+G agarose beads (Beyotime) for 5 h at 4 °C. The beads consisting of immunoprecipitates were collected by centrifugation, followed by washing three times with 1 mL of lysis buffer. For western blot analysis, the harvested protein samples were resuspended in SDS-PAGE loading buffer, boiled at 95 °C for 10 min, subjected to 10%–12% SDS-PAGE, and transferred to polyvinylidene difluoride (PVDF) membranes (Millipore, Burlington, MA, USA). PVDF membranes containing bound proteins were blocked with PBS-Tween containing 5% non-fat dry milk, followed by incubation with the indicated primary and secondary antibodies.

### LC-MS/MS

LC-MS/MS was performed as previously described(Rao *et al*, 2018). HEK293T cells were transfected with pCMV-Tag2B-Flag-IRF3 or empty vector. At 48 h after transfection, the cells were harvested and subjected to the IP experiment with anti-IRF3 antibody. A portion of the protein samples from the IP complex were used to examine IRF3 expression by western blot using antibody against IRF3 and silver staining. LC-MS/MS analysis was performed on a Q Exactive mass spectrometer (Thermo Fisher). MS/MS spectra were searched using the MASCOT engine (Matrix Science) against the Human UniProt sequence database and human database from the NCBI Sequence Read Archive browser.

### ChIP

The cells were crosslinked at 37 °C for 10 min in 1% formaldehyde solution, glycine was added at a final concentration of 125 mM to quench the formaldehyde, and the reaction was terminated at room temperature for 5 min. After washing twice with cold PBS, the cells were collected by centrifugation at 2000 × g for 5 min. Cells at a concentration of approximately 4 × 10^6^ cells/mL were suspended in SDS lysis buffer and kept on ice for 10 min, followed by sonication and subsequent removal of the precipitate by centrifugation. The precipitate was dissolved using 500 μL of 1× ChIP buffer, and 10 μL was removed for the 2% input sample. Anti-Myc or anti-IRF3 antibodies were added to the corresponding sample groups, and IgG protein was added to the negative control group. Then 30 μL ChIP grade G agarose beads were added to each group, followed by incubation for 2 h. The beads were washed thoroughly and centrifuged to collect the precipitate. The precipitate was added to 150 μL ChIP elution buffer and centrifuged to collect the antibodies and IgG in the supernatant, followed by the addition of 6 μl 5M NaCl and 2 μl proteinase K to the supernatant and incubation at 65 °C for 2 h to remove the antibodies from the DNA. The free DNA fragments were purified from the supernatant using DNA purification columns and subjected to RT-PCR.

### ELISA

The IFN-β levels in cell supernatants and mouse sera were measured with ELISA kits purchased from BioLegend (San Diego, CA, USA).

### Determination of viral TCID_50_

The tested virus samples were serially diluted 10 times in DMEM medium. Each serial dilution of the virus was used to inoculate HEK293T cells or RAW264.7 cells, which were cultured at 37 °C in a 5% CO_2_ incubator. The cells were monitored for cytopathic effects (CPEs) twice daily, and the number of CPEs were recorded until the control cells entered senescence. The TCID_50_ values were calculated using the Reed-Muench two-factor method.

### PFU determination

The virus samples were serially diluted 10 times in DMEM medium. HEK293T cells were incubated with diluted virus solutions for 1 h at 37 °C and 5% CO_2_. Unbound viruses were removed and fresh medium was added to the cells, followed by a sustained culture period of 2–3 d at 37 °C and 5% CO_2_. Upon the occurrence of evident cytopathic effects, the cells were incubated with neutral red dye for 2 h and the number of plaques were counted.

### Histopathology analysis

The lungs of wild-type and *Ddx18^+/-^* mice infected with HSV-1 were fixed with paraformaldehyde for 12 h. The fixed tissues were sent to Wuhan Saville Biology Co., Ltd for further paraffin-embedding, sectioning, and hematoxylin-eosin staining in accordance with standard procedures. The results were analyzed by light microscopy.

### Statistical analyses

The results were analyzed for significance by Student t test using GraphPad Prism 6 software (GraphPad Software, Inc.; La Jolla, CA, USA). The difference between groups was considered statistically significant when the p value was < 0.05. All experiments were repeated at least three times.

## Data availability

The data that support the findings of this study are available from the corresponding author on request.

## Acknowledgements

We thank Drs Chunfu Zheng and Zhigao Bu for providing reagents. This work was supported by grants from the National Natural Science Foundation of China (31730095 and 31672569).

## Author contributions

S.X. designed and supervised the study; X.X., M.W., and W.Z. performed most experiments and analyses; P.F., Y.Z., D.W. and L.F. provided reagents and suggestions; X.X., L.F. and S.X. wrote the paper.

## Competing interests

The authors declare no competing financial interests.

